# KBase Research Agent: Automated Multi-Agent Workflow Construction for Reproducible Genome Analysis^*^

**DOI:** 10.64898/2026.06.01.729336

**Authors:** Prachi Gupta, William J. Riehl, Mikaela Cashman, Dylan Chivian, Christopher J. Neely, Shane R. Canon, Robert Cottingham, Chris Henry, Adam P. Arkin, Paramvir S. Dehal

## Abstract

Constructing multi-step bioinformatics workflows, from read quality control through genome assembly to functional annotation, requires expertise in both biology and computational tool selection, creating a bottleneck for scalable and reproducible analysis. We present the KBase Research Agent, a multi-agent system for automating such workflows within the DOE Systems Biology Knowledgebase (KBase). Given a set of sequencing reads and a research objective, the agent constructs an analysis plan grounded in KBase documentation and a Knowledge Graph (KG) of the KBase application catalog, then selects, parameterizes, validates and executes appropriate KBase applications to carry out the workflow. The resulting analysis is preserved as a reproducible KBase Narrative. We evaluate the system’s planning and execution quality against ground truth constructed from reference workflows derived from peer-reviewed Microbiology Resource Announcements. We further apply the agent to 100 previously unanalyzed bacterial isolate genomes from the JGI IMG/M database, where it autonomously performed read quality control, genome assembly, taxonomic classification with GTDB-Tk, and downstream analysis producing annotated genomes, reproducible Narratives, and draft manuscripts without human intervention. Across these experiments, the KBase Research Agent demonstrates the feasibility of domain-grounded, end-to-end scientific workflow automation in a production bioinformatics platform.

## 1 Introduction

The Department of Energy Systems Biology Knowledgebase (KBase) [1] is a platform for biological research and analysis, offering reference data, analysis tools, and a narrative interface for presenting results. Scientists have used KBase for developing many unique workflows, including assembly and annotation of genomes from de novo sequencing, construction of metabolic models of both individual species and microbial communities, and exploration of microbial diversity of novel and distinct ecosystems.

KBase is a growing resource, with a current catalog of apps that make over 300 bioinformatics tools available to the community. These can be brought together, along with narrative text, to perform rich analyses that describe and make testable biological predictions. KBase also provides tutorials and published protocols to help users navigate the system and perform their own experiments over their data.

The platform features a vast and expanding catalog of different bioinformatics tools, and assembling a valid, multi-step analysis pipeline such as for genome assembly and annotation requires expertise. Currently, researchers must manually select, configure, and sequence specific tools from available options, a process that demands interdisciplinary expertise in both biology and computational workflow management. This manual orchestration creates a bottleneck which can lead to inefficiencies and potential reproducibility gaps that hinder the scalability of high-throughput biological research. To bridge this gap, a dedicated KBase research agent is essential. The KBase Research Agent is a novel agentic system designed to automate and streamline complex bioinformatics workflows within the KBase ecosystem. By automating the complex logic of tool selection and workflow generation, such an agent can democratize access to KBase’s powerful high-performance computing resources, ensuring that sophisticated bioinformatics pipelines are streamlined, scalable, and accessible to the broader scientific community.

Advancements in large language models (LLMs) have demonstrated their capability in scientific reasoning, task automation, and knowledge retrieval. These have been used to reason over data and use tools to perform scientific analyses, and assist with interpretation of the results. To achieve robust scientific workflow automation, our work leverages the latest techniques in agentic AI architecture. We move beyond the single-agent paradigm to design a Multi-Agent System (MAS)[2, 3, 4], a collaborative approach that enables groups of LLM-powered agents to coordinate and solve complex tasks by sharing knowledge. Combining the capabilities of individual LLM-based agents, the KBase Research Agent seeks to harness collective intelligence by specializing LLMs into distinct agents, each with unique roles and tools. In this framework, the user goal is achieved through a hierarchical state graph composed of four distinct workflows. A startup workflow initializes the state with metadata, which is then passed to the planning workflow which designs a research plan and gets the approval. Subsequently, the state is passed to the execution workflow which then runs and compiles the results and passes it to the Microbial Resource Announcement (MRA) generation workflow which produces the write up in MRA format. This modular architecture with multiple role based agents that engage in planning, validation and execution mirrors the cooperative nature of human teamwork to solve complex problems that would be beyond the scope of any single agent.

Within this MAS, we leverage the ReAct paradigm to develop task-specific agents that synergize reasoning with KBase specific tool calls to achieve autonomous KBase workflow execution. The central tenet in our system involves prompting the LLM to generate both explicit reasoning traces (thoughts) and task specific tool calls (actions) in an interleaved, iterative manner. This creates a closed-loop system where scientific logic informs the selection of KBase applications, and the execution of those tools grounds the reasoning process in reproducible data objects stored within the KBase Narrative. The agent first generates a reasoning trace to decompose the scientific objective, formulating a valid analysis plan. Based on this detailed step by step plan, the agent generates an action to run a specific KBase app. Together, this hierarchical architecture of ReAct agents within a MAS equipped with the breadth of KBase tools and services enables a powerful system performing scientific analyses, and storing the steps that lead to publishable results While KBase supports a wide variety of analysis types, the present work deliberately focuses on a single but highly representative family of workflows i.e. genome assembly and annotation from microbial isolate reads. This choice reflects both the prevalence of this workflow in KBase usage and the availability of high quality ground-truth Narratives. Throughout the paper we restrict our claims to this workflow family, and we discuss generalization to other KBase analysis types as the next step for future work (Section 7).

This paper introduces the KBase Research Agent, the first LLM based agentic system interfaced with the DOE KBase Narrative environment that translates a user’s research goal and sequencing metadata into an executable, provenance-preserving bioinformatics workflow plan, run the appropriate KBase apps, and draft an MRA compatible results narrative. This hybrid multi-agent architecture couples a LangGraph state machine (for robust lifecycle orchestration and human approval checkpoints) with specialized execution crews and tools. We further contribute a domain specific evaluation framework derived from expert, peer reviewed Narratives to assess both planning validity and execution reliability. The resulting impact is a substantial reduction in the human expertise and time required to generate complete annotated isolate genomes and taxonomy classifications from raw reads, with the added benefit that analyses are automatically captured as reproducible Narratives. This agent provides a scalable path for expanding high quality genomic resources and creates a foundation for extending agentic automation to other KBase workflows.

## Contributions

This initial release of the KBase Research Agent focuses on using isolate genomic reads to produce annotated genomes. These are quality controlled, assembled (if necessary) and annotated by the agent, using the KBase Narrative system. The resulting genomes, along with the pipeline presented in their Narratives, can be published as new genomic resources available to the community, and a draft release can be automatically created from the results. This work makes several contributions to the field of applied scientific AI agents:

1. **Domain-grounded agent infrastructure for KBase:** We develop an agent framework grounded in KBase-specific knowledge sources, including retrieval over KBase documentation, tutorials and knowledge graph (KG) constructed from the KBase app catalog. These resources enable the system to select tools and compose workflows using validated platform-specific knowledge. This hybrid architecture combines two agentic frameworks. At the highest level, we leverage LangGraph [5] for flexible, stateful orchestration of the entire research lifecycle-from planning to execution and reporting. For the execution phase, we instantiate specialized, process-driven CrewAI [6] teams.
2. **Automated KBase app selection, parameterization, and execution:** We develop an agent system that translates research goals and input data into executable KBase workflows by selecting appropriate KBase applications, configuring their required parameters and inputs, and launching analyses automatically. Because these analyses are executed within KBase, the resulting workflows and outputs are preserved as KBase Narratives.
3. **An End-to-End Workflow:** The KBase Research Agent provides a complete solution that automates the full scientific workflow, from interactive user goal-setting and metadata collection, through workflow planning and validation, to automated execution and generation of draft manuscripts.
4. **Domain-Specific Evaluation Framework:** To rigorously validate our system, we develop domain-specific evaluation datasets derived from peer-reviewed scientific narratives. We employ a scalable evaluation protocol using an “LLM-as-a-Judge” methodology to assess the scientific validity of the agent’s planning logic and the reliability of its execution capabilities against expert-curated published ground truth.

## 2 Related Work

### 2.1 LLM-based Agents for Scientific Discovery and Bioinformatics

The deployment of LLM agents to automate complex scientific tasks has emerged as a rapidly growing field. Unlike general-purpose assistants, scientific agents are characterized by their integration with domain-specific tools, structured reasoning capabilities, and access to external knowledge bases. Recent work has also explored agent-based systems for computational and experimental biology. CellAgent [7] applies a multi-agent framework to automated single-cell data analysis, while CRISPR-GPT [8] extends domain-grounded agent design to gene-editing experiment planning and analysis. Together, these efforts highlight a broader shift toward agentic systems that support increasingly complex biological workflows. In chemical sciences, ChemCrow [9] demonstrated the ability of LLM agents to autonomously plan and execute chemical syntheses by integrating 18 expert-designed tools. Similarly, SciAgents [10] applied multi-agent graph reasoning to materials science, revealing hidden interdisciplinary relationships through ontological knowledge graphs. Within the bioinformatics domain, several systems have recently been proposed to address the high barrier to entry for computational genomics. AutoBA [11] introduced an autonomous agent capable of generating analysis plans for multi-omics data with minimal user input. BioMANIA [12] focused on an inquiry-based assistant for processing omics data, while BioAgents [13] and BioMaster [4] leveraged multi-agent architectures to decompose complex genomic tasks into manageable code generation steps and CellAgent[7] uses a plan,execute and evaluate style multi-agent architecture for automated single-cell data analysis. Our work extends this paradigm by integrating agentic capabilities directly with the KBase Narrative infrastructure, employing a robust LangGraph state machine to ensure that the generated workflows are not only executable but also strictly adhere to the reproducibility standards required for peer-reviewed publication. These reproducibility standards are also evident through the use of the KBase Narrative, itself designed around reproducibility.

### 2.2 Evolution of Agent Architectures

The architecture of modern AI agents has evolved significantly, progressing from traditional single-agent systems [14] to more complex, collaborative multi-agent frameworks [2]. While powerful, the cognitive load on a single agent has led to the increasing adoption of MAS, which distribute intelligence and specialize roles to solve problems of higher complexity. MAS architectures are often characterized by their structure (e.g., centralized, decentralized) and their collaboration strategies (e.g., role-based). Prominent frameworks such as AutoGen [15] and MetaGPT [16] often employ role-based workflows, which are effective but can be rigid.

The foundational work on agentic reasoning established paradigms such as Chain-of-Thought (CoT) prompting [17], which elicits reasoning by having the models generate intermediate steps. This was advanced by the ReAct framework [18], which synergizes reasoning and acting by prompting the LLM to interleave verbal thoughts with actions (tool use), grounding the agent’s internal reasoning in external environmental feedback.

Our work builds on these principles by adopting a hybrid architecture. We use CrewAI [6] for its strengths in managing process-driven, role-based execution crews, while leveraging LangGraph[5] for its more flexible, graph-based state machine that enables complex, cyclical orchestration with clear points for human-in-the-loop validation.

### 2.3 Agent Evaluation Methods

The rapid advancement of agent capabilities necessitates a parallel evolution in evaluation methodologies to ensure reliability and measure progress. Early efforts produced comprehensive, multi-dimensional benchmarks like AgentBench [19], which assesses general agent capabilities across a diverse set of eight environments, including operating systems, databases, and web browsing. Another foundational benchmark, GAIA[20], introduced a “human-simple, AI-hard” philosophy, focusing on real-world questions that require robust tool use.

More recently, the field has seen a rise in domain-specific benchmarks that provide more targeted evaluations. Examples include VisualWebArena [21] for multimodal web navigation and, of particular relevance to our work, ScienceAgentBench [19] for evaluating agents on data-driven scientific discovery tasks. Furthermore, as architectures have shifted towards collaborative systems, benchmarks like MultiAgentBench [22] have been developed to explicitly measure the dynamics of coordination and competition.

## 3 System Architecture and Methods

To develop a flexible, modular, and controllable system for automating bioinformatics analyses, we designed a multi-agent framework using LangGraph, a stateful agent framework built on top of the LangChain ecosystem. LangGraph allows for the definition of workflows as directed graphs, where each node represents a specific function or behavior and the edges represent transitions based on conditions or outcomes. This architecture is particularly suited for complex workflows that require iterative planning, conditional execution, and Human-in-the-loop (HITL) validation.

Each workflow in our system is implemented as a LangGraph graph, which operates over a persistent shared state. The agent is composed of multiple modular nodes, each encapsulating a specific task such as planning, execution, or summarizing. The LangGraph framework provides fine-grained control over the flow of logic between these nodes, enabling the agent to make decisions based on intermediate results, user input, or feedback from external tools.

This LangGraph-based system supports loopbacks (e.g., for re-planning or re-execution), parallel branches (e.g., for modular execution of independent steps), and asynchronous execution, making it well-suited for automating multi-step scientific workflows such as genome annotation, metagenome assembly, and report generation.

The agent workflow operates through four main phases, each of which are separate agentic graph workflows that make use of tools that either work directly with user data in KBase, or query against KBase documentation and app capabilities. These phases are metadata collection, planning, execution, and draft manuscript generation.

**Figure 1:**
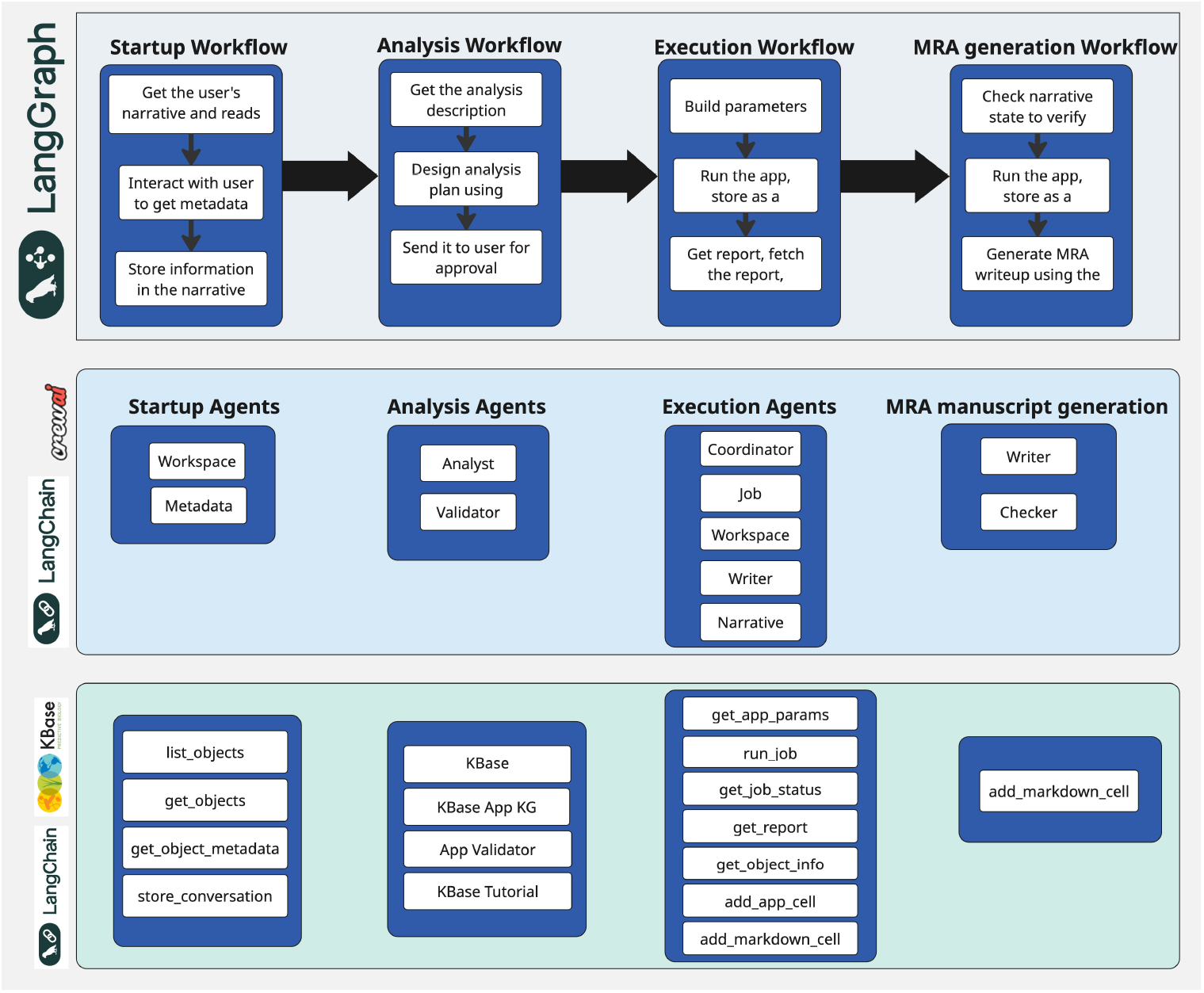
Hierarchical multi-agent architecture of the KBase Research Agent. The system is organized into three vertically integrated layers: **(Top) The Orchestration Layer** (implemented in LangGraph) manages the high-level state transitions and control flow across four lifecycle stages: Startup, Analysis Planning, Execution, and MRA Generation. It enforces logic for human-in-the-loop approvals and error recovery. **(Middle) The Agent Layer** consists of specialized LangChain agents (e.g., *Analyst, Validator, Coordinator*) that are dynamically instantiated to handle specific cognitive tasks within each workflow stage. **(Bottom) The Tool Layer** provides the agents with grounded capabilities, including wrappers for the KBase API (e.g., `run_job, list_objects`) and RAG tools for accessing KBase documentation and the App Knowledge Graph.

**Figure 2:**
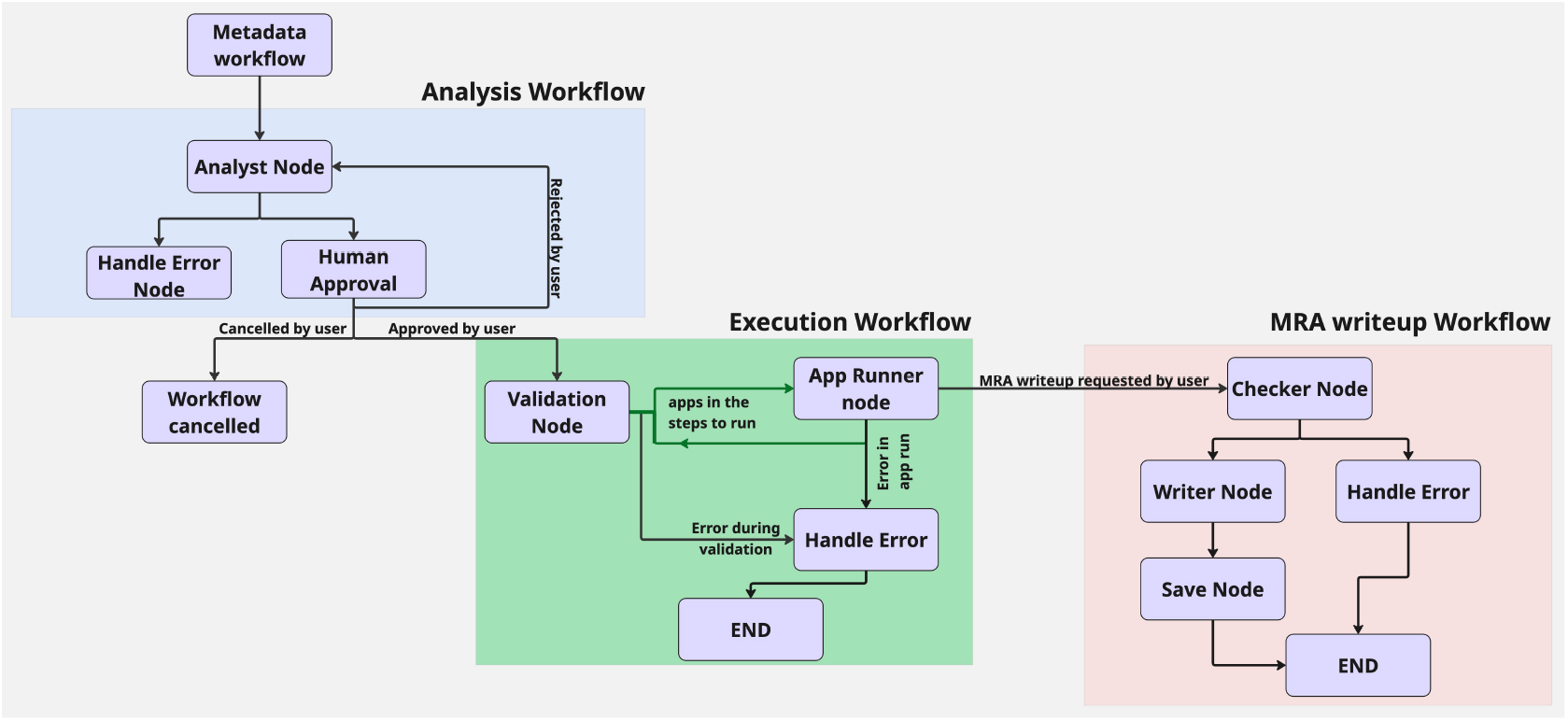
The hierarchical state graph architecture of the KBase Research Agent implemented in LangGraph. The workflow is composed of three coupled graphs that manage the life cycle of a scientific task: (1) The **Analysis Workflow** (blue) synthesizes a research plan and enforces a HITL checkpoint, requiring explicit user approval or feedback before proceeding. (2) The **Execution Workflow** (green) manages the interface with the KBase Narrative. It features a self-correction loop where the *Validation* and *Handle Error* nodes can iteratively refine parameters to resolve runtime failures without human intervention. (3) The **MRA Writeup Workflow** (red) triggers post-execution to compile results into a draft manuscript, employing a *Checker Node* to verify the consistency of the generated text against the analysis data.

### 3.1 Metadata Collection Workflow

The goal of the metadata collection workflow is to extract sample metadata from user-provided data. It first takes data from the user’s selected KBase data object, and starts a conversation with the user, focused on refining any needs the user might have or other metadata that might be missing but would be otherwise useful for choosing the right set of apps to run to achieve analysis goals.

The user is prompted to enter any known information about the data or experiments that generated it, with a focus on determining data types (i.e. single-end or paired-end reads), and specifics about it such as read length, insert size, and sequencing depth. It then asks the user about any preferred bioinformatics tools used for processing. This information is then collated and digested into a form that can be passed to the planning workflow, and is then stored in the Narrative.

### 3.2 Planning Workflow

The planning workflow is initiated after metadata collection and is orchestrated by LLM powered planning and validation nodes, whose role is to convert user intent and metadata into a structured, step-by-step bioinformatics analysis plan.

The planning workflow uses a ReAct agent that uses Chain of Thought to elicit reasoning with predefined tools. These tools use Retrieval-Augmented Generation (RAG) [23] to ensure accurate plan generation. This retrieved context is dynamically injected into the prompt presented to the language model, grounding the generation in known practices and tool capabilities.

**Figure 3:**
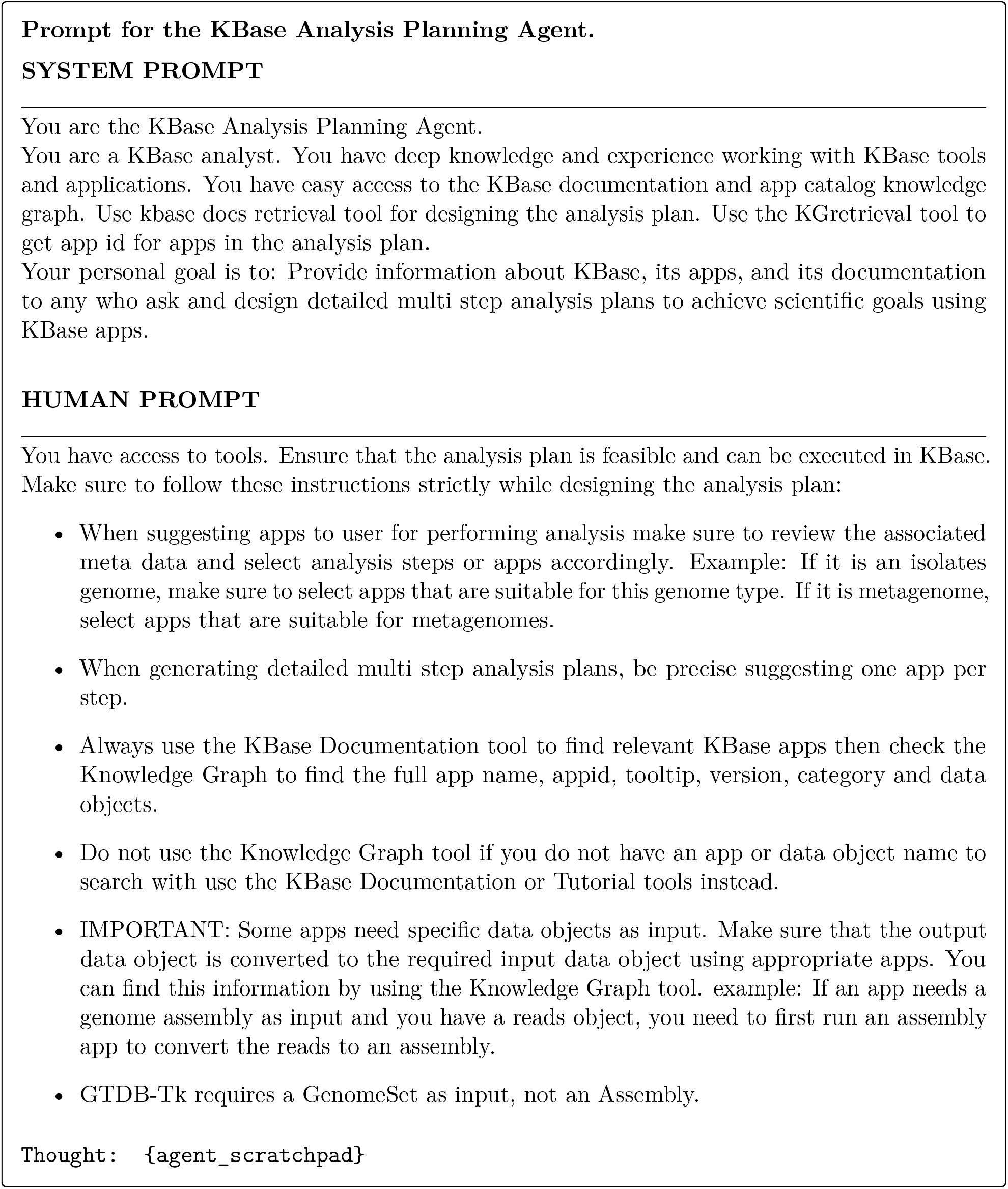
Prompt template used by the KBase Analysis Planning Agent.

The prompt includes:

- A natural language description of the sample and user goals
- Planning directions and MRA heuristics
- Details about the tools available to the agent
- Constraints or preferences

As part of the planning process, the system leverages a Knowledge Graph (KG) tool to validate and refine app selections proposed by the planning agent. The tool retrieves the precise app ID for each app which are stored in the shared state alongside the workflow plan, ensuring that execution nodes can directly call the correct tool without further disambiguation.

The generated plan is passed to a human approval node, which creates a condensed version suitable for human review. If HITL approval is enabled, the agent pauses execution and waits for confirmation or revision. The system supports plan editing or full re-planning through loopbacks in the graph structure.

#### 3.2.1 Planning Agent Tools

To ensure accuracy, domain relevance, and robustness, the planning agent uses LLM powered tools that perform RAG. Specifically, before invoking the language model, the agent retrieves relevant domain knowledge from structured and unstructured sources, including:

- Internal KBase documentation on supported bioinformatics tools
- Prior curated workflows and analysis examples from published tutorial narratives
- A KBase KG constructed from app catalog containing information about the KBase applications

##### Documentation and Tutorial RAG

The KBase documentation and tutorial RAG system operates by first indexing documentation content into a vector database during the setup phase. Pages are chunked into semantically meaningful segments, and each chunk is converted into dense vector embeddings using a transformer-based encoder. Our retrieval pipeline is implemented with a Chroma vector store built over KBase documentation pages and tutorial materials. Documents are split using a recursive character-based text splitter with a chunk size of 1000 characters and an overlap of 20 characters, which provides enough local context for tool descriptions and usage instructions while preserving retrieval granularity. We generate embeddings with the nomic-embed-text-v1.5 model at 768 dimensions and persist the resulting vector indexes separately for each corpus. At inference time, the agent retrieves relevant chunks from these indexed resources to ground planning and tool selection in KBase-specific documentation rather than relying only on parametric model knowledge. During the analysis workflow, when agents need to design analysis plans or understand KBase apps, they generate query embeddings and perform similarity searches against the indexed documentation. The most relevant chunks are retrieved based on cosine similarity scores, then provided as context to the language model along with the original query.

##### Knowledge Graph Construction

The KBase KG is constructed from the KBase app catalog. It encodes structured relationships between KBase applications, their functional categories, accepted input and output data types, and app descriptions. The knowledge graph is constructed in Neo4j by creating nodes representing KBase entities (apps and data objects) and establishing typed relationships that capture their semantic connections.

Retrieval operates through a two-stage process: first, all app names are fetched from the graph, then TF-IDF vectorization is applied to both the query entity and the retrieved app names to generate feature vectors. Cosine similarity is calculated between the query vector and each app vector, with apps ranked by similarity score in descending order. The highest-ranking match is then used to query Neo4j for comprehensive node information.

The user approved plan is then passed down to the validation node to validate the coherence of the proposed plan (e.g., are the input/output objects in the various steps consistent). Once the plan is validated under set heuristics it is passed to the execution workflow.

### 3.3 Execution Workflow

Following approval of the analysis plan, the execution workflow performs each step in sequence, interfacing with the KBase platform to run the corresponding applications and capture outputs. Once the HITL review confirms the analysis plan, the system transitions into the execution phase, where computational jobs are run on the KBase platform.

The workflow follows a cyclical validation-execution pattern where graph execution begins at the validation node with an initial state containing the complete analysis plan from the planning phase. For each step in the validation cycle, the validator confirms input object availability and parameter completeness, with invalid states triggering error handling while valid states proceed to execution.

This phase is managed by a Job Crew, implemented using the CrewAI framework, and orchestrated within the overall LangGraph HITL workflow. CrewAI enables multi-agent collaboration during execution, with specialized agents responsible for job preparation, submission, monitoring, and result handling.

#### 3.3.1 Execution Agents

The job execution node instantiates a CrewAI multi-agent crew to handle app-specific parameter construction, job submission, monitoring, report retrieval, and analysis. This crew employs specialized agents:

- **CoordinatorAgent:** Constructs app-specific parameters using dynamic Pydantic models generated from KBase app specifications
- **JobAgent:** Submits jobs to the KBase Execution Engine and monitors completion status
- **WorkspaceAgent:** Retrieves output objects and reports from the KBase Workspace
- **WriterAgent:** Interprets job reports (HTML, JSON, or plain text) and generates biological insights
- **NarrativeAgent:** Persists analysis summaries as markdown cells in the target Narrative

During the execution cycle, the CrewAI crew orchestrates a five-task pipeline that begins with parameter construction using app specifications and input object metadata, followed by job submission and monitoring via the KBase Execution Engine, then report retrieval from the Workspace using the job’s report data, subsequently performing report interpretation and biological analysis, and finally persisting the analysis as markdown in the Narrative.

Upon task completion, the node updates the workflow state by removing the completed step from steps to run, appending it to completed steps, and recording any created objects with their unique identifiers and names. The router then evaluates the updated state and either cycles back to validation for the next step or terminates execution if the queue is exhausted.

### 3.4 Draft Publication Generation Workflow

After all workflow steps are completed, an optional draft publication generation workflow is performed which follows the formatting for a MRA. The Agent is prompted with the MRA format, and the content of the Narrative that was previously assembled from the app runs. It generates text conforming to the format by listing the bioinformatics tools used, the experimental goals, data used, and the generated genome. The result leaves space for the user to put in relevant figures and references.

### 3.5 Automatic Pipeline

Each of the above steps have also been assembled into an automatic pipeline that removes the HITL components. This allows a user to upload external data files to their KBase staging area, or select an available data object in KBase, then run the script which automatically generates Narratives, genomes, and reports for each input file.

### 3.6 Computing environment

The LLM components of the KBase Research Agent were run through CBORG on commercial cloud infrastructure. KBase analysis jobs were executed on KBase’s internal compute cluster, which consists of 37 nodes with approximately 3,300 CPUs and 38 TB of RAM. This cluster includes 28 smaller 64-CPU nodes and 9 larger 168-CPU nodes, primarily running CentOS 7, with one node running AlmaLinux 9. The workflows evaluated in this study were executed sequentially rather than as parallelized job graphs. However, the platform supports running multiple workflows in parallel for a single user.

## 4 Evaluations

In this section, we discuss our evaluation datasets, models and methodology. Evaluating the quality of agent generated scientific workflows presents a significant challenge. Standard metrics for natural language generation, such as BLEU or ROUGE, are insufficient as they do not capture the semantic correctness, executability, or scientific validity of a complex, multi-step plan [24]. Given the focus on reproducible methods and robust workflows for scientific computing, a rigorous evaluation methodology is paramount. We adopt a dual-evaluation framework that integrates scalable, automated assessment with targeted expert review. First, we employ an “LLM-as-a-Judge” (LLM-a-J) evaluator [25], which scores workflow plans using a structured rubric designed to assess correctness, coherence, and executability. Second, domain experts conduct a qualitative review of both (1) a sample of the generated workflows and (2) the reasoning and scoring logic produced by the judge model. This review of LLM judge by domain experts is a best practice established by other high-stakes scientific benchmarks, such as the unified, multi-dimensional AstroVisBench in astronomy [26].

### 4.1 Data Collection

To rigorously evaluate the agent’s ability to generate complex, real-world scientific workflows, we constructed a ground truth (GT) benchmark dataset from published, peer-reviewed Microbiology Resource Announcements (MRA) papers. We curated our datasets by collecting KBase Narratives associated with these MRA publications. The narratives are then processed to obtain the goal and steps that were used for analysis making this the GT for planning evaluations. This process ensures that each GT workflow is not only scientifically valid but has also passed peer review.

Each data point in our benchmark dataset consists of a complete, multi-step bioinformatics workflow. A typical example is a genome assembly and annotation pipeline, which involves a specific sequence of “Apps” (KBase’s term for tools). For instance, a workflow might proceed from read quality control (e.g., using FastQC), to assembly (e.g., using SPAdes), and finally to functional annotation (e.g., using Prokka or RAST). The ground truth for each task is the precise sequence of these applications and their configurations as documented in the published Narrative.

To capture the output of our own agent-based system for evaluation, we utilize LangSmith [27] for comprehensive tracing and logging.

### 4.2 LLMs used for Evaluation

For the workflow planning evaluation, agent configurations were tested based on three popular LLMs: “Claude Sonnet High” is Anthropic’s Claude 4.5 Sonnet model with thinking enabled, “Claude Sonnet” is Anthropic’s Claude 4.5 Sonnet model, and “GPT-5” is OpenAI’s GPT-5 model. This selection allows for a comparative analysis of plan quality which is presented in Section 5. In the execution phase of the KBase Research Agent the agents construct app parameters, submit jobs to the KBase Execution Engine, monitors job status, and interprets reports. The execution phase requires highly structured, low-variance behavior: the LLM must generate consistent parameter dictionaries, follow strict input/output schema, and avoid hallucinated tool names or unsupported parameter fields. For the execution workflow, we use GPT-4.1 since it has demonstrated exceptionally stable tool-calling behavior, producing reproducible parameter structures across repeated runs.

### 4.3 LLM as a Judge Evaluation

To ensure that the evaluation method is scalable, we use an LLM judge to evaluate the analysis and execution workflow. This automated approach assesses the quality of a generated workflow by invoking a separate LLM tasked with scoring the output based on a detailed, rubric-based prompt.

We integrate our LLM-as-a-Judge evaluator within LangSmith’s evaluation framework, following an OpenEvals [28] style setup in which the judge model is given both the agent generated workflow and the corresponding GT workflow extracted from the MRA Narratives. The judge compares the generated plan or execution state directly against this reference GT and assigns a rubric-based score. This pipeline produces both detailed traces and structured evaluation records, enabling systematic comparison between agent outputs and published GT workflows.

The evaluation rubric provided to the judge model is structured around several key dimensions critical to scientific workflows:

- **Workflow Validity and Task Completion:** Assesses whether the agent generated plan is valid and successfully completes the stated scientific task.
- **Scientific Correctness:** Evaluates the factual accuracy of the plan’s components and the correct use of domain-specific tools and methods.
- **Relevance and Coherence:** Measures whether the generated plan directly addresses all parts of the scientific objective and follows a logical, coherent structure

#### Evaluation prompt and rubric

To ensure consistent, rubric-driven scoring across datasets and model configurations, we designed two structured evaluation prompts: one for the planning workflow and one for the execution workflow. These prompts provide explicit scoring criteria, define acceptable variation in workflow structure, and guide the judge model to produce reliable, interpretable assessments. The planning evaluation prompt (PLANNING_CUSTOM_PROMPT) instructs the judge model to act as an expert bioinformatics workflow evaluator with specialization in KBase analysis plans. The prompt incorporates a detailed rubric that emphasizes five dimensions of workflow quality:

- **Step Coverage**: Whether the generated plan includes all essential stages of a genomic workflow.
- **App Selection**: Whether selected KBase apps are appropriate, with explicit allowance for functionally equivalent alternatives.
- **Logical Flow**: Whether workflow steps follow a scientifically sound order and maintain correct input–output dependencies.
- **Completeness**: Inclusion of quality control, validation, and other required preprocessing or postprocessing steps.
- **Accuracy**: Whether step descriptions correctly state their purpose.

The rubric (Table 1) defines six scoring tiers, enabling continuous scoring while maintaining qualitative interpretability. The prompt directs the judge model to compare the generated plan to the ground-truth workflow step-by-step, identify missing or incorrect steps, assess functional equivalence in app substitutions, and provide a brief rationale for the assigned score. This structured design ensures rubric compliance and mitigates over-penalization for scientifically acceptable variation in app choice.

**Table 1:** Plan quality scoring rubric used in our evaluation. Scores are assigned on a continuous scale from 0.0 to 1.0 based on step coverage, app appropriateness, and workflow ordering.

| Rating | Score Range | Criteria |
| --- | --- | --- |
| Excellent | 0.90–1.00 | Plan includes all essential steps with appropriate (or functionally equivalent) KBase apps and correct ordering. Minor differences in app choices are acceptable if they are functionally equivalent. |
| Good | 0.70–0.89 | Plan covers most essential steps with mostly appropriate apps. May have minor gaps in coverage or slight ordering issues, but the overall workflow is sound. |
| Adequate | 0.50–0.69 | Plan includes core steps but misses some important quality control or validation steps. Apps are generally appropriate but may include some suboptimal selections. |
| Poor | 0.30–0.49 | Plan is incomplete, missing several important steps. Some inappropriate app selections. Workflow order may have issues. Descriptions may contain inaccuracies. |
| Very Poor | 0.10–0.29 | Plan is severely incomplete or incorrect. Many inappropriate apps and major gaps in workflow logic. |
| Incorrect | 0.00 | Plan is fundamentally flawed, missing most essential steps, or does not utilize KBase apps appropriately. |

The execution scoring prompt (`EXECUTION_EVALUATION_PROMPT`) evaluates the workflow execution state rather than workflow structure. The judge model is asked to assess whether each step executed successfully by examining fields such as `step_result, job_status, completed_steps`, and `error`. The rubric prioritizes:

- **Successful job completion**: job_status = completed and absence of errors.
- **Absence of failure signals**: any non-null value in error indicates step-level or workflow-level failure.
- **Progress through workflow**: higher scores require progression through assembly, annotation, and validation steps.

The scoring tiers reflect how far the workflow progressed before encountering errors. For example, workflows that complete all major steps receive *Excellent* scores (0.9–1.0), whereas those that fail prior to assembly receive *Very Poor* or *Incorrect* scores (0.0–0.29).

#### Transparency and bias mitigation

To reduce ambiguity and potential evaluator bias, the judge model is always given both the agent generated workflow and the corresponding ground-truth workflow extracted from the MRA Narrative, and is instructed to score only with respect to the explicit rubric criteria. The rubric explicitly allows functionally equivalent KBase apps (e.g., SPAdes versus megahit for assembly, RAST versus Prokka for annotation), so that scientifically valid substitutions are not penalized.

## 5 Experimental Results and Analysis

In this section, we present evaluation results assessing the performance of the KBase Research Agent across both the planning and execution stages of the agentic workflow. Our evaluation uses three datasets derived from peer-reviewed Microbiology Resource Announcements (MRA) Narratives as mentioned in Section 4.1 and compares multiple LLM configurations for planning and execution reliability.

### 5.1 Planning workflow evaluation

The planning workflow was evaluated using three foundation models: Claude Sonnet 4.5 (standard), Claude Sonnet 4.5 with thinking enabled, and GPT-5. As summarized in Table 2 all models achieved consistently high scores under the LLM-as-a-Judge rubric, indicating strong capability in decomposing scientific objectives into coherent, executable multi-step workflows. Claude Sonnet 4.5 (thinking enabled) achieved the highest mean score (*0*.*906* on the EXP_MRA_0084523 dataset), suggesting that the additional reasoning traces improve planning reliability. Across all datasets, the narrow score differences among the three models indicate that all models are broadly capable of producing high-quality, scientifically grounded workflow plans. Across repeated planning runs, the agent produced stable, high-quality workflows when requests completed successfully, with plans typically remaining in the Good–Excellent range under our rubric (consistent end-to-end structure with only minor app/order variation).The planning runs that did not include a post trimming quality check were penalized and got a “Good” rating rather than excellent. Similar penalization was observed for missing “Annotate and Distill Assemblies with DRAM” step. The main failure cases we observed in the high-throughput setting were transient provider rate-limit events, which interrupted the analyst agent before it could return a valid structured plan. Downstream, this surfaced as a workflow-state validation error (*steps*_*to*_*run* = *N one*), i.e., a cascading failure from rate limiting. We evaluate the run-to-run bias for *EXP* _*MRA*_0084523 dataset with Claude Sonnet and Claude Sonnet thinking. We observed low run-to-run variability in planning quality over 5 repetitions per organism with both models. For Claude Sonnet, mean rubric scores ranged from adequate to good, trending good, with very small variances (0.002–0.008). The most stable cases were Bacillus strain SC127 (variance 0.002) and Bacillus strain SC123 (variance 0.003), while the highest variability was still modest for Bacillus velezensis SC119 (variance 0.008) and Priestia megaterium SC120 (variance 0.007). For Claude Sonnet with thinking, mean rubric scores were consistently high (Good to near-Excellent), and run-to-run variance remained low overall (0.003 for four of five datasets). The only notably higher variability was for Bacillus strain SC123 (variance 0.010), while the other organisms showed tightly clustered performance: SC127 (0.003), B. pumilus SC124 (0.003), B. velezensis SC119 (0.0033), and Priestia megaterium SC120 (0.003). Overall, these results indicate the planner is highly reproducible across repeated runs, with only minor score fluctuations between repetitions.

**Table 2:**
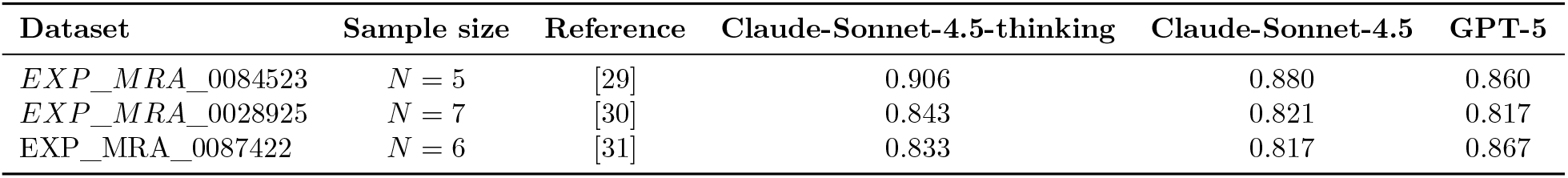
Detailed breakdown of evaluation benchmarks.

| Dataset | Sample size | Reference | Claude-Sonnet-4.5-thinking | Claude-Sonnet-4.5 | GPT-5 |
| --- | --- | --- | --- | --- | --- |
| *EXP_MRA_0084523* | $N = 5$ | [29] | 0.906 | 0.880 | 0.860 |
| *EXP_MRA_0028925* | $N = 7$ | [30] | 0.843 | 0.821 | 0.817 |
| *EXP_MRA_0087422* | $N = 6$ | [31] | 0.833 | 0.817 | 0.867 |

### 5.2 Execution workflow evaluation

The execution workflow evaluation measures the system’s ability to correctly execute the plan passed to it by the planning agent. GPT-4.1 is used for the execution of all datasets and results were assessed using the LLM-as-a-Judge scoring rubric described in Section 4.3, which evaluates the final state for `job_status, step_result, completed_steps`, and the presence or absence of errors. Table 3 summarizes the execution performance across the three benchmark datasets. These results indicate that the KBase Research Agent is capable of reliably executing multi-step workflows, correctly handling the transition between various steps, and producing outputs consistent with the published Narrative results.

**Table 3:** Detailed breakdown of execution evaluation benchmarks.

| Dataset | Sample size | Reference | Mean score | Execution time per genome (hours) |
| --- | --- | --- | --- | --- |
| EXP_MRA_0084523 | $N = 5$ | [29] | 0.840 | 1.024 |
| EXP_MRA_0028925 | $N = 7$ | [30] | 0.943 | 1.311 |
| EXP_MRA_0087422 | $N = 6$ | [31] | 0.867 | 1.739 |

## 6 Executing the KBase Research Agent workflow to perform Genome Assembly and Annotation

### 6.1 Overview

The automatic pipeline was applied to a set of 100 publicly available paired-end reads sequenced from isolate bacterial genomes. This workflow was used to operate on these reads to execute comprehensive genome assembly and annotation. Each analysis produced a separate KBase Narrative, including a draft of a MRA. All narratives include the analysis apps that were run, the inputs and parameters used, the outputs produced, and a detailed analysis of each output with its biological significance.

### 6.2 Data Preparation

Genomic reads were identified in the Joint Genome Institute’s (JGI) Integrated Microbial Genomes and Microbiomes (IMG/M) database [32] using the following criteria:

- Only bacterial reads were chosen.
- Reads were in “Permanent Draft” status, denoting that though they were incomplete and unpublished, they were not expected to be used in any further analyses by JGI.
- All data must be public.
- They must not have an associated Pubmed or Genbank id, denoting that they are not part of published genomes.

This resulted in 4533 unique sequencing project ids. Reads files were downloaded from 100 different genomes. These were all uploaded to the KBase data staging service and imported into a single Narrative as individual paired-end reads objects, and made publicly available. [33].

### 6.3 Procedure

The automatic KBase Research Agent pipeline was run on each of the JGI read sets. First, individual Narratives were created for each read set, and those reads were copied into the Narrative as the starting data object. Object metadata was automatically generated by KBase’s data import app. This included details about the reads: number of reads, their mean and standard deviation length; minimum, maximum, mean, and standard deviation quality values; total base count, and total read count. The sequencing technology field was kept as “Unknown” during the import process, as that information was either missing or inconsistent in the downloaded data packet. However, the agents seemed able to infer likely machine sources based on the metadata.

Once the metadata was loaded, an automated prompt was used for the agent to interpret the metadata and create an abstract cell in the Narrative with a summary of the reads and goals for the workflow.

**Figure 4:**
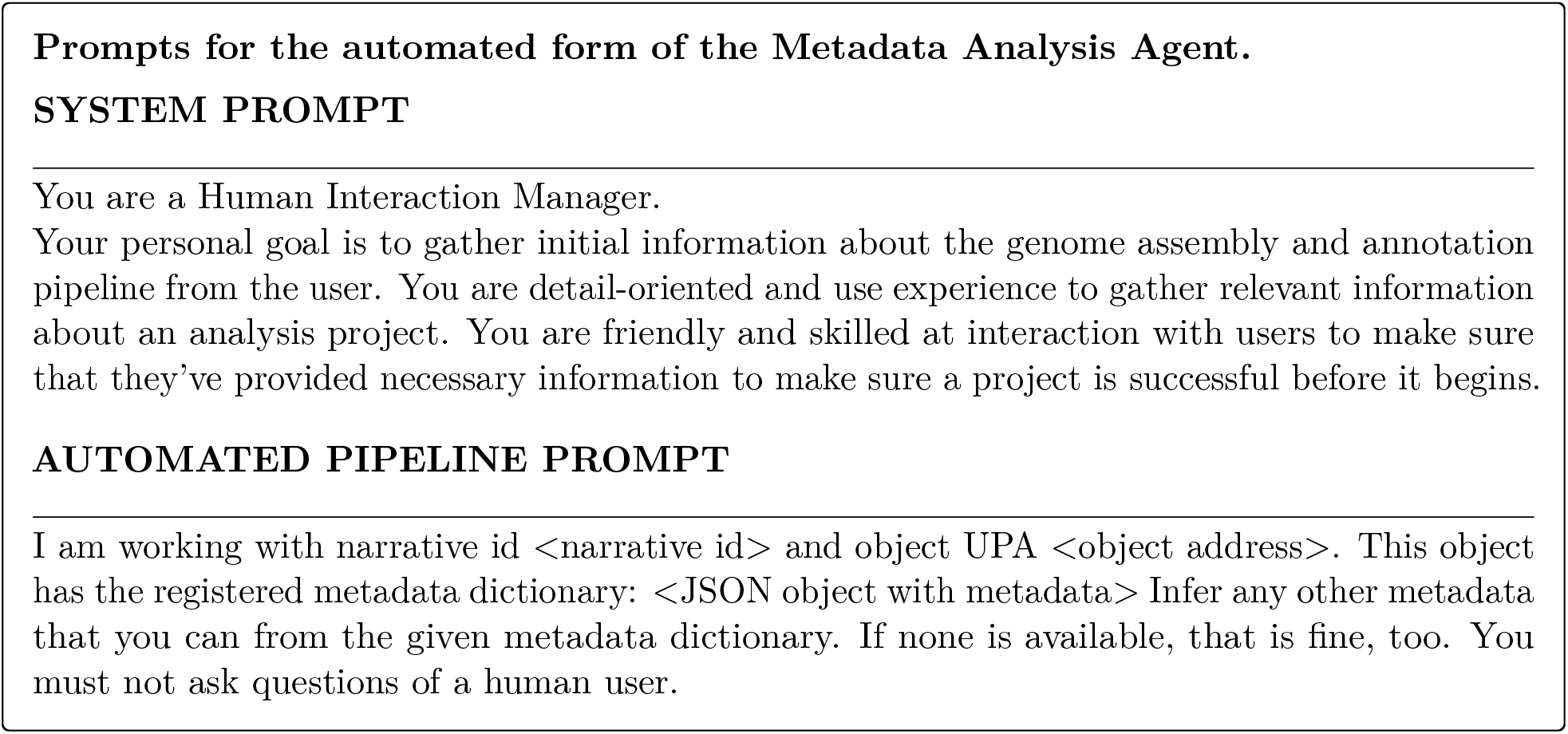
Prompt template used by the Metadata Analysis Agent.

### 6.4 Results

One hundred Narratives were created from the reads sets downloaded from JGI, each generated from the automated pipeline. These results were evaluated based on their final classification of the annotated genome into a genetic clade using GTDB-Tk [34]. Original clades from the sequencing project were provided in the metadata downloaded from JGI, these were used as a point of comparison. Due to the nature of the sequencing project, many of these clades were labeled as members of the Actinomycetota phylum, while the results from GTDB-Tk were more specific.

Some of the reads remained unclassified by GTDB-Tk. In these cases, the report showed that the resulting genome was either too fragmented or incomplete for a full phylogenetic analysis. While assembly and annotation was still possible using the available tools, the reads data used was not of sufficient quantity, due possibly to a lack of coverage depth. Other cases (labeled “Missing results”) failed to utilize the full pipeline. The reads used in these cases were either dismissed by the agents as being unusable after initial quality control, or because of unrecoverable errors that happened during the pipeline app runs. See Table 4 for details.

**Table 4:** Narrative LLM Agent Prediction Comparison Summary. GTDB-Tk predictions compared against IMG-derived genome classifications, stratified by taxonomic match level.

| Category | Count | Percentage |
| --- | --- | --- |
| Total matches (any taxonomic level) | 74 | 74.0% |
| Exact match (genus + species) | 15 | 15.0% |
| Partial match (same genus) | 16 | 16.0% |
| Phylum-level match | 45 | 45.0% |
| Mismatches | 6 | 6.0% |
| Unclassified | 13 | 13.0% |
| Missing results | 5 | 5.0% |

These results show that although the sets of reads chosen from the IMG database had variable quality, the KBase Research Agent was still able to work with them, and in many cases produce an annotated and taxonomically classified genome that may be of use to the research community. See [35] for full data results and links to all agent-generated Narratives.

## 7 Discussion

The development of the KBase Research Agent represents a significant step toward democratizing access to high-performance computing resources for bioinformatics. By integrating Large Language Models with the structured, reproducible environment of the KBase platform, we addressed the bottleneck of manual workflow orchestration.

The results presented in Sections 5 and 6.4 demonstrate that the KBase Research Agent is capable of autonomously generating and executing scientifically valid bioinformatics workflows, marking a significant step toward scalable and reproducible analysis within the KBase ecosystem. By combining a hierarchical LangGraph-based architecture with specialized execution crews and retrieval-based planning agents, the system bridges a gap between manual expert-driven workflow construction and automated, end-to-end bioinformatics analysis. The evaluation on 100 isolate genomes (Section 6.4) further illustrates the scalability of the approach. The agent consistently produced complete genome assembly and annotations, each backed by a fully reproducible KBase Narrative, without human intervention. This demonstrates that the system can operate not only on curated benchmarks, but also on realistic, heterogeneous sequencing datasets, supporting high-throughput applications where manual curation would be prohibitively slow. There was also a demonstration of cases where the agent judged that completing the workflow would not be useful, based on the data used, and that workflow was stopped, avoiding extraneous resource usage.

A limitation of the present paper is that our current agentic framework and its evaluation focuses on assembly and annotation workflows for isolate microbial reads. Although this is a common and scientifically important workflow family in KBase, it does not cover the full diversity of analysis types supported by the platform, such as metagenome assembly, pangenome analysis, or metabolic modeling. Extending the system to these workflow classes will require expanding the agent’s tool repertoire, enriching its workflow-specific planning prompts, and broadening the retrieval corpus used to ground action selection. For example, pangenome analysis would require support for comparative genomics tools while metabolic modeling would require access to model-building, gapfilling, and flux analysis tools together with prompts that encode biological constraints, media conditions, and model-quality checks. Extending our evaluations and benchmark datasets to include multiple, representative workflow classes is an important next step. We view the current work as a first demonstration that end-to-end agentic workflows can reproduce bioinformatics pipelines in KBase, providing a foundation for broader and more diverse automated scientific analyses.

We will continue to refine the KBase research agent in several ways. (1) We aim to enhance autonomous debugging and error-handling capabilities, enabling the system to detect, diagnose, and recover from execution failures without human intervention. (2) We are expanding the agent’s applicability to a broader range of bioinformatics workflows, allowing it to leverage the full breadth of KBase tools while maintaining strict reproducibility standards. (3) We plan to further improve and rigorously evaluate agent performance across diverse scientific and biological domains, ensuring that the system remains robust, generalizable, and scientifically trustworthy. Overall, the KBase Research Agent demonstrates that end to end bioinformatics workflow automation that requires substantial human expertise can be achieved through the use of a MAS. The approach presented here establishes the foundation for scalable, reproducible, and democratized access to computational biology analysis within KBase.

## 8 Code Availability

All code and configuration needed to reproduce our experiments are publicly available in the KBase Research Agent repository [36].

## 9 Acknowledgments

This work was supported as part of the BER Genomic Science Program. The DOE Systems Biology Knowledgebase (KBase) is funded by the U.S. Department of Energy, Office of Science, Office of Biological and Environmental Research under Award Numbers DE-AC02-05CH11231, DE-AC02-06CH11357, DE-AC05-00OR22725, and DE-SC0012704. This research used the CBorg AI platform and resources provided by the IT Division at the Lawrence Berkeley National Laboratory (Supported by the Director, Office of Science, Office of Basic Energy Sciences, of the U.S. Department of Energy under Contract No. DE-AC02-05CH11231)

## Notes

https://github.com/kbaseincubator/narrative_llm_agent

### Competing Interest Statement

The authors have declared no competing interest.

## References

[1] Adam P. Arkin, Robert W. Cottingham, Christopher S. Henry, Nomi L. Harris, Rick L. Stevens, Sergei Maslov, Doreen Ware, Fernando Perez, Shane Canon, Michael W. Sneddon, Matthew L. Henderson, William J. Riehl, Dan Murphy-Olson, Stephen Y. Chan, Roy T. Kamimura, Sunita Kumari, Meghan M. Drake, Thomas S. Brettin, Elizabeth M. Glass, Dylan Chivian, Dan Gunter, David J. Weston, Benjamin H. Allen, and Jason Baumohl. KBase: The United States Department of Energy Systems Biology Knowledgebase. Nature Biotechnology, 36(7):566–569, jul 2018.

[2] Khanh-Tung Tran, Dung Dao, Minh-Duong Nguyen, Quoc-Viet Pham, Barry O’Sullivan, and Hoang D. Nguyen. Multi-agent collaboration mechanisms: A survey of llms, 2025.

[3] Shervin Minaee, Tomas Mikolov, Narjes Nikzad, Meysam Chenaghlu, Richard Socher, Xavier Amatriain, and Jianfeng Gao. Large language models: A survey, 2025.

[4] H. Su, W. Long, and Y. Zhang. BioMaster: Multi-agent System for Automated Bioinformatics Analysis Workflow. bioRxiv, 2025.

[5] LangChain, Inc. LangGraph. https://github.com/langchain-ai/langgraph, 2024.

[6] CrewAI Inc. CrewAI: Framework for orchestrating role-playing, autonomous AI agents. https://github.com/crewAIInc/crewAI, 2023.

[7] Yihang Xiao, Jinyi Liu, Yan Zheng, Xiaohan Xie, Jianye Hao, Mingzhi Li, Ruitao Wang, Fei Ni, Yuxiao Li, Jintian Luo, Shaoqing Jiao, and Jiajie Peng. Cellagent: An llm-driven multi-agent framework for automated single-cell data analysis. arXiv preprint arXiv:2407.09811, 2024.

[8] Yuanhao Qu, Kaixuan Huang, Ming Yin, Kanghong Zhan, Dyllan Liu, D. Yin, Henry C. Cousins, William A. Johnson, Xiaotong Wang, Mihir Shah, Russ B. Altman, Denny Zhou, Mengdi Wang, and Le Cong. Crispr-gpt for agentic automation of gene-editing experiments. Nature Biomedical Engineering, 2025.

[9] Andres M. Bran, Sam Cox, Oliver Schilter, Carlo Baldassari, Andrew D. White, and Philippe Schwaller. ChemCrow: Augmenting large-language models with chemistry tools. Nature Machine Intelligence, 6(5):525–535, 2024.

[10] Alireza Ghafarollahi and Markus J Buehler. Sciagents: Automating scientific discovery through multi-agent intelligent graph reasoning. arXiv preprint arXiv:2409.05556, 2024.

[11] J. Zhou, B. Zhang, G. Li, X. Chen, H. Li, X. Xu, S. Chen, W. He, C. Xu, L. Liu, and X. Gao. An AI Agent for Fully Automated Multi-Omic Analyses. Advanced Science, 11(e2407094), 2024.

[12] Sameen Sohail, Haoyang Li, Xiaopeng Xu, Juexiao Zhou, and Xin Gao. Biomania: Biomedical agent for navigation, inquiry, and analysis. arXiv preprint arXiv:2312.03604, 2023.

[13] Nikita Mehandru, Amanda K Hall, Olesya Melnichenko, Yulia Dubinina, Daniel Tsirulnikov, David Bamman, Ahmed Alaa, Scott Saponas, and Venkat S Malladi. Bioagents: Democratizing bioinformatics analysis with multi-agent systems. arXiv preprint arXiv:2501.06314, 2025.

[14] Michael Wooldridge and Nicholas R Jennings. Intelligent agents: Theory and practice. The knowledge engineering review, 10(2):115–152, 1995.

[15] Qingyun Wu, Gagan Bansal, Jieyu Zhang, Yiran Wu, Shaokun Zhang, Erkang Zhu, Beibin Li, Li Jiang, Xiaoyun Zhang, and Chi Wang. AutoGen: Enabling Next-Gen LLM Applications via Multi-Agent Conversation Framework, 2023.

[16] Sirui Hong, Mingchen Zhuge, Jonathan Chen, Xiawu Zheng, Yuheng Cheng, Jinlin Wang, Ceyao Zhang, Zili Wang, and others. MetaGPT: Meta Programming for a Multi-Agent Collaborative Framework. In The Twelfth International Conference on Learning Representations (ICLR), 2023.

[17] Jason Wei, Xuezhi Wang, Dale Schuurmans, Maarten Bosma, Fei Xia, Ed Chi, Quoc V Le, Denny Zhou, et al. Chain-of-thought prompting elicits reasoning in large language models. Advances in neural information processing systems, 35:24824–24837, 2022.

[18] Shunyu Yao, Jeffrey Zhao, Dian Yu, Nan Du, Izhak Shafran, Karthik Narasimhan, and Yuan Cao. ReAct: Synergizing Reasoning and Acting in Language Models. In International Conference on Learning Representations (ICLR), 2023.

[19] Xiao Liu, Hao Yu, Hanchen Zhang, Yifan Xu, Xuanyu Lei, Hangliang Ding, Kejuan Yang, Shudan Zhang, and others. AgentBench: Evaluating LLMs as Agents. In International Conference on Learning Representations (ICLR), 2024.

[20] Grégoire Mialon, Clémentine Fourrier, Craig Swift, Thomas Wolf, Yann LeCun, and Thomas Scialom. GAIA: a benchmark for General AI Assistants, 2023.

[21] Jing Yu Koh, Robert Lo, Lawrence Jang, Vikram Duvvur, Ming Chong Lim, Po-Yu Huang, Graham Neubig, Shuyan Zhou, Ruslan Salakhutdinov, and Daniel Fried. VisualWebArena: Evaluating Multimodal Agents on Realistic Visual Web Tasks. In Proceedings of the 62nd Annual Meeting of the Association for Computational Linguistics (Volume 1: Long Papers), pages 881–905, Bangkok, Thailand, aug 2024. Association for Computational Linguistics.

[22] Kunlun Zhu and others. MultiAgentBench: Evaluating the Collaboration and Competition of LLM agents, 2025.

[23] Patrick Lewis, Ethan Perez, Aleksandra Piktus, Fabio Petroni, Vladimir Karpukhin, Naman Goyal, Heinrich Küttler, Mike Lewis, Wen-tau Yih, Tim Rocktäschel, et al. Retrieval-augmented generation for knowledge-intensive nlp tasks. Advances in neural information processing systems, 33:9459–9474, 2020.

[24] Yang Chen, Shaoxiong Ji, et al. Rethinking evaluation metrics for long-form scientific text generation. In Proceedings of the 2023 Conference on Empirical Methods in Natural Language Processing, 2023.

[25] Helia Hashemi, Jason Eisner, Corby Rosset, Benjamin Van Durme, and Chris Kedzie. LLM-Rubric: A Multidimensional, Calibrated Approach to Automated Evaluation of Natural Language Texts. In Proceedings of the 62nd Annual Meeting of the Association for Computational Linguistics (Volume 1: Long Papers), pages 13806–13834, Bangkok, Thailand, 2024. Association for Computational Linguistics.

[26] Sebastian Joseph et al. AstroVisBench: A Code Benchmark for Scientific Computing and Visualization in Astronomy. arXiv preprint, 2505.20538, 2025. Accepted at NeurIPS 2025 Datasets & Benchmarks Track.

[27] LangChain, Inc. Langsmith: Observability and evaluation for llm applications. https://smith.langchain.com, 2024. Accessed: 2025-12-12.

[28] LangChain, Inc. Openevals: Readymade evaluators for llm applications. https://github.com/langchain-ai/openevals, 2024. Accessed: 2025-12-12.

[29] Anna L McLoon, Prince Ackaah Asante, Thomas Anderson, Kellyanne Cahill, Delana Cochrane, Keira Cohen, Jaylene German, Christian M Hrubes, Isabella LaCroix, Killian McNamee, et al. Five draft genome assemblies from bacillaceae isolated from a degraded wetland environment. Microbiology Resource Announcements, 13(2):e00845–23, 2024.

[30] Anna L McLoon, Julia M Barker, Gillian M Churan, Anthony Cucca, Lillian Gardner, Justin Gejo, Francesca Gerbasi, Michaela Higgins, Lillian Kronau, Jalin Le, et al. Draft genome assemblies from seven bacillaceae isolates from woodland soil. Microbiology Resource Announcements, 14(6):e00289–25, 2025.

[31] Anna L McLoon, Thomas T Awad, Molly F Bogardus, Meredith G Buono, Kaitlyn A Devine, Rebecca M Draper, Brianna Femenella, Hannah M Gallagher, Laura A Morelock, Mishal Razi, et al. Draft genome sequences for 6 isolates of endospore-forming class bacilli species isolated from soil from a suburban, wooded, developed space. Microbiology Resource Announcements, 11(11):e00874–22, 2022.

[32] I-Min A Chen, Ken Chu, Krishnaveni Palaniappan, Anna Ratner, Jinghua Huang, Marcel Huntemann, Patrick Hajek, Stephan J Ritter, Cody Webb, Dongying Wu, Neha J Varghese, T B K Reddy, Supratim Mukherjee, Galina Ovchinnikova, Matt Nolan, Rekha Seshadri, Simon Roux, Axel Visel, Tanja Woyke, Emiley A Eloe-Fadrosh, Nikos C Kyrpides, and Natalia N Ivanova. The IMG/M data management and analysis system v.7: content updates and new features. Nucleic Acids Res., 51(D1):D723–D732, January 2023.

[33] Prachi Gupta, William J. Riehl, and Paramvir S. Dehal. Jgi unpublished reads. https://narrative.kbase.us/narrative/239038, 2025. Accessed: 2025-12-12.

[34] Pierre-Alain Chaumeil, Aaron J Mussig, Philip Hugenholtz, and Donovan H Parks. GTDB-Tk v2: memory friendly classification with the genome taxonomy database. Bioinformatics, 38(23):5315–5316, November 2022.

[35] Prachi Gupta, William J. Riehl, and Paramvir S. Dehal. Jgi reads processing results. https://github.com/kbaseincubator/narrative_llm_agent/blob/main/jgi_data_results/img_llm_annotations_with_gtdb.tsv, 2025. Accessed: 2025-12-12.

[36] Prachi Gupta, William J. Riehl, and Paramvir S. Dehal. Kbase research agent. https://github.com/kbaseincubator/narrative_llm_agent, 2025. Accessed: 2025-12-12.

